# Growing microalga *Neochloris oleoabundans* in rocking and floating plastic bag photobioreactors

**DOI:** 10.64898/2026.02.18.702949

**Authors:** Sergei A. Markov, Samantha L. Childs, Jared K. Averitt, Richard A. Johansen

## Abstract

This paper evaluated and compared the relative microalgal biomass accumulation of rocking, floating, and stationary bag photobioreactors. Microalga *Neochloris oleoabundans* was grown in these photobioreactors in batch mode for 24 days under illumination. The 50 L plastic bags (cell suspension volume 25 L) were placed on the surface of a rocking platform, an artificial pond or a stationary platform. In the pond, waves were generated by electrical fans which shake and mix microalgal cells within the plastic bags. The bags were supplied with 5% CO_2_ in air under elevated pressure inside of the bags. The rocking bag method significantly increased biomass yields to approximately 3-4 g • L^-1^, as compared to 0.16 g • L^-1^ in the floating photobioreactor and only 0.03 g • L^-1^ in the stationary type photobioreactor.

## 1. Introduction

Current industrial production of photosynthetic microalgae (microscopic algae) for practical purposes is accomplished in artificial ponds of several hundred hectares in size [1, 2]. Maintaining these systems endures several challenges such as low volumetric productivity, water loss and difficulty preventing contamination [3, 4]. Additional issues with the pond approach are temperature control and land usage. The demand to overcome these limits and to promote microalgal biotechnology to the level of bacterial cultivation has led to the development of photobioreactors that efficiently utilize solar energy and monitor culture purity [5]. Photobioreactors consist of closed systems made of transparent materials in which microalgae are cultivated under control conditions and exposed to light [6,7,8]. The perfect photobioreactor should be low-cost, with high volumetric and areal productivity, and energy efficient.

A vast amount of work to design and optimize different photobioreactor systems for microalgal cultivation has been carried out and reviewed [6, 7, 9, 10]. The most common type of inexpensive photobioreactors for microalgal cultivation, described in scientific literature, are tubular and flat panel (tank) photobioreactors made of glass or plastic. The tubes or tanks are generally thin to allow deep light penetration. Microalgae in photobioreactors also need water, CO_2_ and mineral nutrients for their growth. The main problem in photobioreactor design is to achieve the efficient supply of CO_2_ (carbon source for algae) and light (energy source for algae) because light intensity and CO_2_ concentrations decrease with distance from the surface of a photobioreactor [11]. These challenges encouraged us to design our experiments with plastic bag photobioreactors where algal suspension mixing was provided by continuous rocking of the photobioreactor, and CO_2_ was supplied under elevated pressures.

Here, we developed, tested and compared three types of photobioreactors in relation to microalgal biomass accumulation: rocking-type photobioreactors with plastic bags placed on a rocking platform, floating bag photobioreactors and stationary plastic bag photobioreactors. In 2009, we acquired a series of BIOSTAT CultiBag bioreactors from Sartorius Stedim Biotech which are made of a rocking platform and a 50 L plastic bag (cell suspension working volume is 25 L) equipped with gas flow sensors. These bioreactors (rocking-type) were used to grow microalgae with external light source [12,13].

Photobioreactors of floating type [14, 15, 16, 17] have been known at least since 90s. One floating type photobioreactor was also developed by NASA (OMEGA project) and tested outdoor but data were not published in scientific literature. The floating bag photobioreactor has many advantages, it requires minimal land space, algal cell suspension can be mixed by waves and is easily cooled in water during hot days. The rocking-type photobioreactors [12, 13, 18, 19], on a contrary, received little attention so far. The rocking-type photobioreactors may not be as practical as the floating type for growing microalgae on industrial scale for some algal applications such as biodiesel generation. However, they still can be used in laboratory settings for study of microalgal growth and biofuel generation or for manufacturing of some highly priced products. They also can be used as simulation models for floating type photobioreactors. Apart from us, several other researchers used rocking type photobioreactors for growing diatoms [18] and cyanobacteria [19].

Our experimental microalga was *Neochloris oleoabundans* which is one of candidates for biodiesel generation, because it is rich in oil [4, 20, 21] and can be utilized in pharmaceutical and cosmetic industries. So given these factors, it was very important for us to find the optimal conditions for *N. oleoabundans* cultivation in photobioreactors.

## 2. Materials and Methods

### 2.1. Microalga

The microalga *Neochloris oleoabundans* (Neo) UTEX #1185 bacteria-free strain was obtained from University of Texas (Austin) Culture Collection. It is a freshwater green alga from *Chlorophyceae* class. Microalga was grown on a tris-acetate-phosphate medium – TAP medium [22]. The alga was also evaluated using Allen-Arnon medium [23] with similar results.

Prior to using the photobioreactor, we grew *N. oleoabundans* in 250-ml Erlenmeyer flasks with cell suspension volume of 100 ml at pH 7. Continuous light was provided by cool white fluorescent lamps (20 μmol • m^-2^ • s^-1^ on the surface of the culture). The cells were harvested at OD_665_ ∼ 0.54 corresponding to the chlorophyll concentration of ∼ 0.05 μg • ml^-1^, and transferred to a photobioreactor (four flasks per 25 L photobioreactor volume).

### 2.2. Photobioreactor description

#### 2.2.1. Rocking bag photobioreactor

BIOSTAT CultiBag RM 20/50 Basic bioreactors from Sartorius Stedim Biotech (Goettingen, Germany), which were originally developed for animal cell culturing, were used in our experiments (Figure 1). The BIOSTAT CultiBag bioreactors are made of a rocker, rocking platform and a 50 L plastic bag (with cell suspension working volume of 25 L) equipped with temperature sensors, gas flow control and adjustable bag rocking speed. The bag size was 95 cm x 50 cm. The rocker is fully automated and operated by a front LCD display with a cyclical menu structure. The rocker also has an aeration flow rate controller. We added the external light source to these bioreactors - cool white fluorescent lamps. Algal cells were illuminated continuously except for our experiments in green house where we used natural light-dark cycle. Temperature sensors of CultiBag bioreactors were not utilized in our experiments.

**Figure 1.**
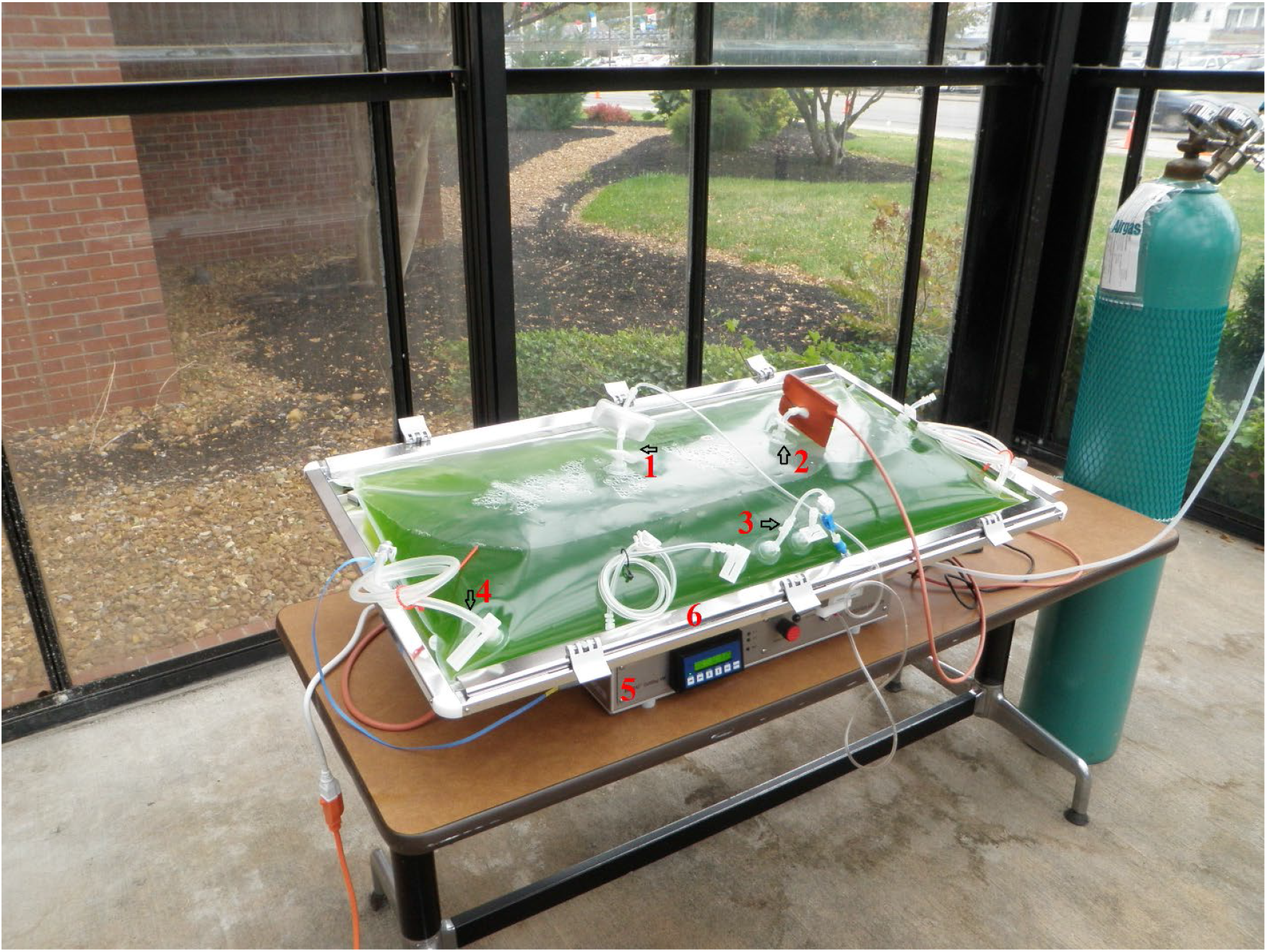
BIOSTAT CultiBag bioreactor with a rocking platform and a 50 L plastic bag in green house under natural illumination. 1) Inlet with sterile filter for 5% CO_2_/air mix. 2) Air outlet with sterile filter, filter heater, and pressure relief valve. 3) Luer-septum for sampling. 4) Inlet/tubing for cell inoculation. 5) Rocker. 6) Bag holder.

#### 2.2.2. Floating bag photobioreactor

We used an artificial pond (above ground portable swimming pool 15’ x 42’) and placed 50 L plastic bags (with cell suspension working volume of 25 L) from Sartorius Stedim Biotech with microalgal cells on the surface of water. The bag type/size was exactly the same as for our rocking bag photobioreactor (Figure 1). We also used an aeration flow rate controller from BIOSTAT CultiBag bioreactor to run the floating bag photobioreactor. We applied the electrical fan (Powermax Electrical Co., LTD., China) to generate waves for our pond in order to mix algal suspension inside of our bags.

#### 2.2.3. Stationary bag photobioreactor

The 50L plastic bags (with cell suspension working volume of 25 L) from Sartorius Stedim Biotech containing microalga *N. oleoabundans* were used as our stationary photobioreactor. The bag type/size was the same as for our rocking bag photobioreactor (Figure 1). We used an aeration flow rate controller from BIOSTAT CultiBag bioreactor to run our stationary bag photobioreactor. This photobioreactor was used as a control for rocking and floating bag photobioreactors.

For all photobioreactors (except in a greenhouse) continuous light was provided by cool white fluorescent lamps (170 - 180 μmol • m^-2^ • s^-1^ on the surface of the culture). The photobioreactor bags were pre-sterilized by Sartorius Stedim Biotech Company. They were inoculated with algal cells under sterile conditions inside our Laminar Flow cabinet. The bags were supplied with gas mixture of 5% CO_2_ and air (with higher gas concentration compared to the ambient conditions). We created an elevated pressure of CO_2_ in the gas phase inside of the bags (exit valve was open when the gas pressure reached a specific point) which helped us to increase the transfer of CO_2_ into microalgal cells. The pH was in the range of 6.74 - 7.74. The gas flow was in the rage of 30-250 mL/min (we started at 30 mL/min and ended with 250 mL/min). The photobioreactor temperature was at the range of 26.6^0^C - 28^0^C.

### 2.3. Cell growth evaluation

The cell growth was evaluated over time in terms of optical density (OD) at 665 nm and chlorophyll content at 665 nm using Spectronic 20 (Spectronic Instruments, Fitchburg, WI) and nanodrop spectrophotometer (NanoDrop Technologies, Wilmington, DE) for extra control. The final optical density value was the average number from three operations of each photobioreactor – rocking, floating and stationary) – three biological replicates. In addition, samples for optical density measurements of algal suspension were taken three times each day at the exact same time (three measurement replicates) and were averaged for each photobioreactor run, thus the total number of repetitions for each photobioreactor was 9. The greenhouse photobioreactor was an exception by only being operated one time, so the total number of optical measurement repetitions for each day was 3.

### 2.4. Chlorophyll content

Chlorophyll content was determined by the method of Talling and Driver (24).

### 2.5. Biomass estimation

Algal biomass from photobioreactor bags was harvested after 24 days during a stationary phase of microalgal culture and concentrated through sedimentation with subsequent drying under 70^°^C in a thermostat. Each photobioreactor (rocking, floating and stationary photobioreactor) was run three times and the final dried biomass value was the average number from these three operations.

## 3. Results

### 3.1. Rocking bag photobioreactor under continuous artificial illumination

Initially, we tested the conditions for microalgal growth in rocking bag photobioreactor, which represent a model system for our floating type photobioreactor. The photobioreactor was tested in a batch mode for 24 days. Figure 2 shows algal cell growth in terms of optical density (OD) in the photobioreactor. The amount of dry algal biomass harvested from 50 L (25 L liquid phase) photobioreactor bags operated for 24 days was on average 73 g, which represents a total biomass yield of approximately 3 g per L.

**Figure 2.**
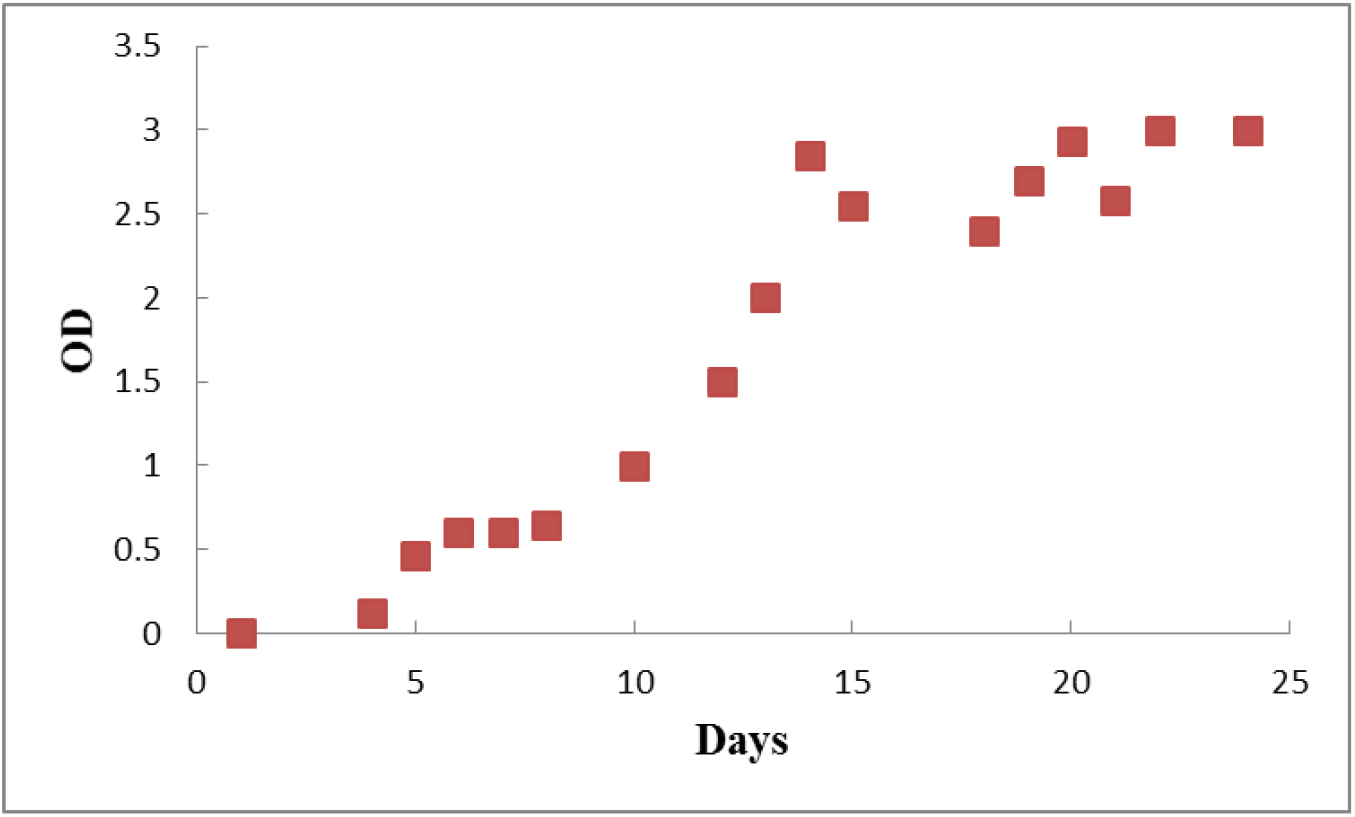
Optical density of *N. oleoabundans* grown under artificial illumination (170 - 180 μmol • m^-2^ • s^-1^ on the surface of the culture) in a rocking bag photobioreactor. Data are shown as mean, n=9.

### 3.2. Rocking bag photobioreactor under natural illumination in greenhouse

The photobioreactor was tested in a batch mode for 24 days in greenhouse during autumn months under natural illumination (Figure 1). Figure 3 shows algal cell growth in terms of optical density (OD).

**Figure 3.**
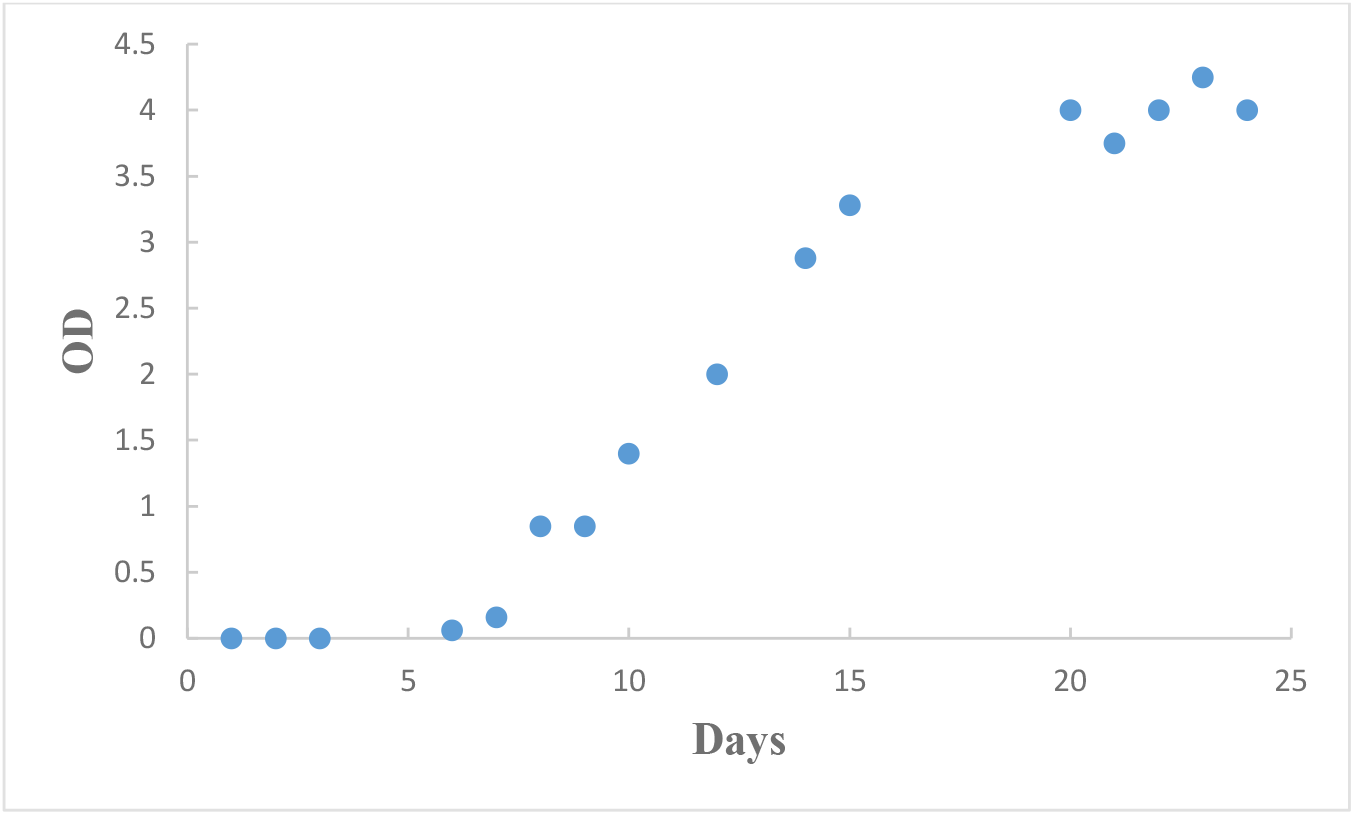
Optical density of *N. oleoabundans* grown under natural illumination in greenhouse in a rocking bag photobioreactor. Data are shown as mean, n=3.

The light intensity was measured twice a day at 12:30 pm and 4:30 pm. Light intensity fluctuated significantly (Figure 4). The amount of dry algal biomass harvested from 50 L (25 L liquid phase) photobioreactor bag operated for 24 days was 96 g, with a biomass yield of approximately 4 g per L.

**Figure 4.**
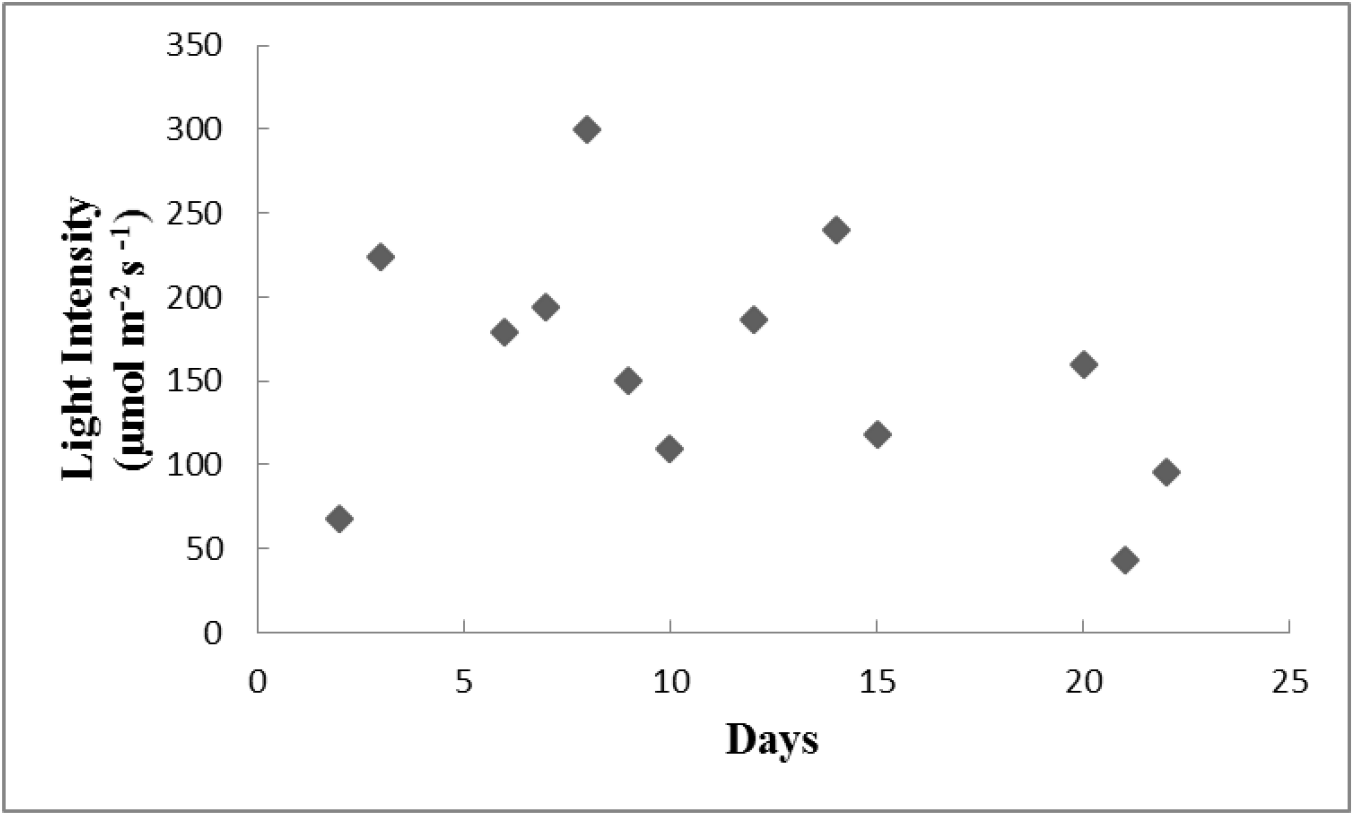
Light intensity on the photobioreactor surface in a greenhouse in October/November at 12:30 pm (Clarksville, TN).

### 3.3. Floating bag photobioreactor under continuous artificial illumination

This photobioreactor was also tested in a batch mode for 24 days. The optical density was evaluated periodically (Figure 5). The amount of dry algal biomass harvested from 50 L (25 L liquid phase) photobioreactor bags operated for 24 days was on average 4 g, with a biomass yield of 0.16 g per L.

**Figure 5.**
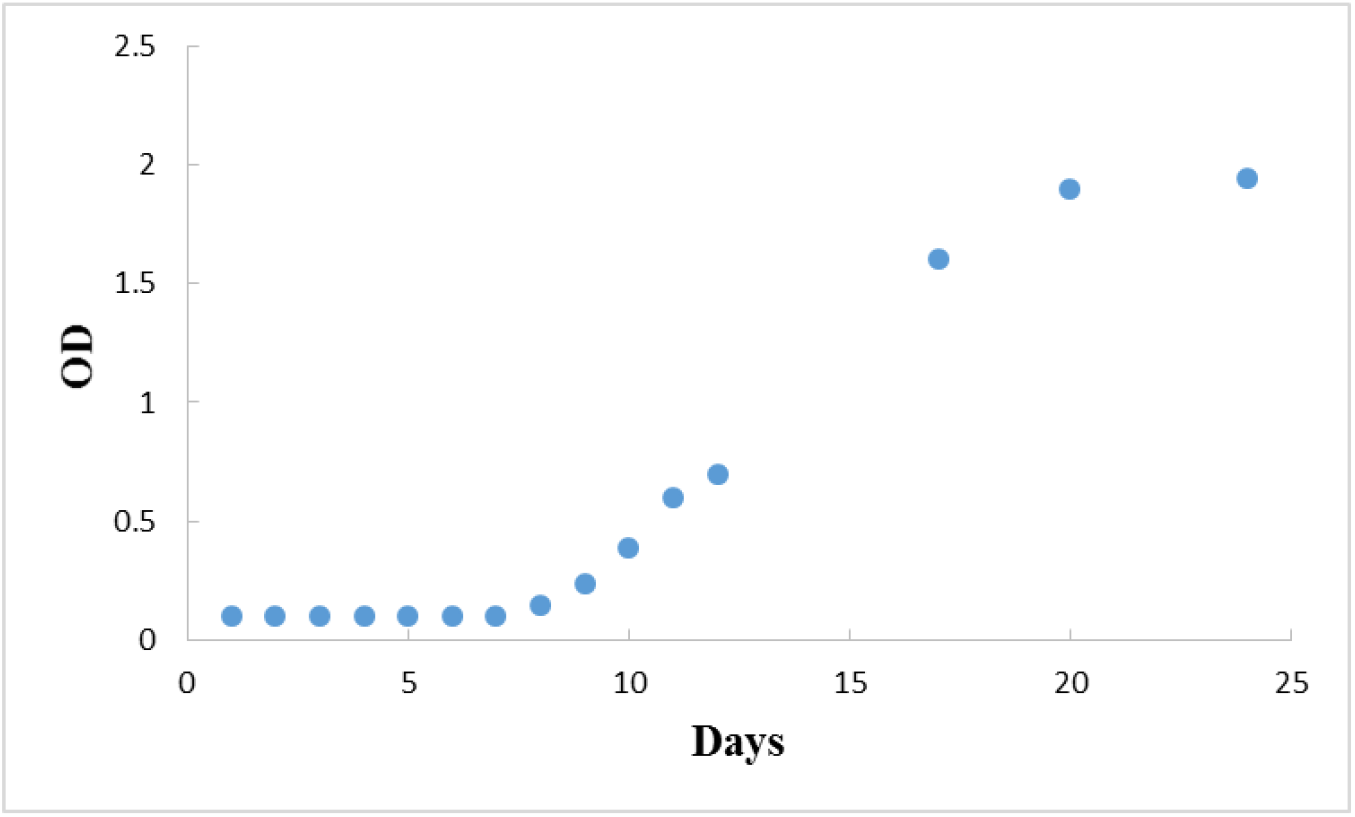
Optical density of *N. oleoabundans* as a function of time in a floating bag photobioreactor under artificial illumination (170 - 180 μmol • m^-2^ • s^-1^ on the surface of the culture). Data are shown as mean, n=9.

### 3.4. Stationary bag photobioreactor under continuous artificial illumination

The photobioreactor was tested in a batch mode for 24 days. The optical density in this photobioreactor did not change dramatically (Figure 6). The amount of dry algal biomass harvested from 50 L (25 L liquid phase) photobioreactor operated for 24 days was on average 0.8 g, with a biomass yield of 0.03 g per L.

**Figure 6.**
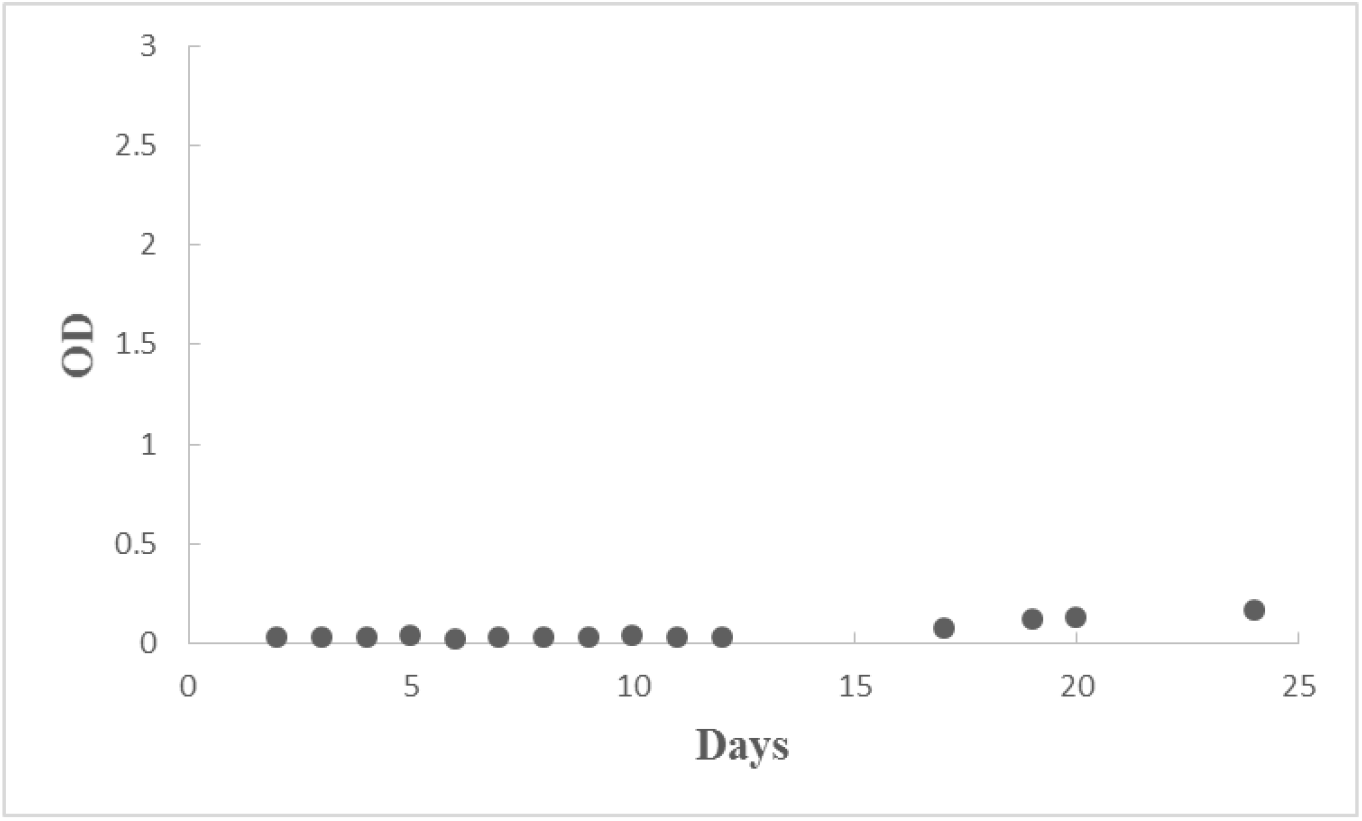
Optical density of *N. oleoabundans* as a function of time in a stationary bag photobioreactor (control) under artificial illumination (170 - 180 μmol • m^-2^ • s^-1^ on the surface of the culture). Data are shown as mean, n=9.

## 4. Discussion

In this study the rocking and floating bag photobioreactors were tested for growth of microalga *N. oleoabundans*. The results surpassed our anticipations by acquiring higher yields in rocking bag photobioreactors (approximately 3-4 g per L) when compared to yield data in floating and stationary bag photobioreactors. Yield data in rocking bag photobioreactors were about 22 times higher than the floating bag and more than 100 times higher compared to stationary bag photobioreactors. These results were possible through mixing the microalgal suspension in the photobioreactor, which allows for more effective light illumination and CO_2_ supply. Optical densities of mixed cultures were on average 15-25 times higher compared to the non-mixing cultures.

Given this data, we believe it is possible to achieve even higher yield by further manipulating CO_2_ supply and light absorption. Our data support this notion: we had higher optical density values if we grew our microalgae under natural illumination (average light intensity was higher compared to light intensity we used for running photobioreactors indoor). The chlorophyll concentrations were also greater under natural illumination (results not shown).

Many photobioreactors were designed and constructed in the past specifically to increase light and CO_2_ supply to maximize photosynthetic capabilities of microalgae. In fact, the main purpose for the novel photobioreactor designs is to increase microalgal growth by manipulating light absorption, CO_2_ supply and other factors. Ideally, light and CO_2_ should reach every single algal cell [25]. One effective approach is the effective mixing of cell suspension using rocking (or floating) photobioreactor, which allows the microalgae to capture more light and, as well as transferring more CO_2_ to individual cells [10]. Mixing of microalgal suspension also helps to reduce the over-exposure of upper cell layers in photobioreactors to higher illumination which can inhibit algal photosynthesis [26].

Our floating bag photobioreactors were very effective for microalgal growth but not as effective as rocking-type photobioreactors presented here. Rocking bag photobioreactors were successfully used by other researchers to study growth and CO_2_ consumption by microalgae. Cires *et al*. [18] were able to increase up to 70% CO_2_ fixation rate of cyanobacterium *Anabaena siamensis* grown in a rocking type photobioreactor compared to bubbled suspension batch cultures. Unfortunately, as we mentioned before, rocking bag photobioreactors are not practical for industrial large-scale applications, but they still could be used in laboratory settings to study microalgae.

## Acknowledgements

This work was supported by National Science Foundation (USA), award # 0821571 (2008) and Austin Peay State University Summer Faculty Research Program. We thank Ms. L. Beiermann for valuable assistance with the experimental work.

## The Statement of Conflicts, Informed Consent, Human/Animal Rights

No conflicts of interest, informed consent, human or animal rights applicable.

## An Author Agreement/Declaration

All authors have made substantial contributions to all sections of this manuscript. Dr. Markov takes full responsibility for the integrity of the work, from inception to finished article.

